# Covalent Attachment of Horseradish Peroxidase to Single-Walled Carbon Nanotubes for Hydrogen Peroxide Detection

**DOI:** 10.1101/2023.12.14.571773

**Authors:** Francis Ledesma, Shoichi Nishitani, Francis J. Cunningham, Joshua D. Hubbard, Dabin Yim, Alison Lui, Linda Chio, Aishwarya Murali, Markita P. Landry

## Abstract

Single-walled carbon nanotubes (SWCNTs) are desirable nanoparticles for sensing biological analytes due to their photostability and intrinsic near-infrared fluorescence. Previous strategies for generating SWCNT nanosensors have leveraged nonspecific adsorption of sensing modalities to the hydrophobic SWCNT surface that often require engineering new molecular recognition elements. An attractive alternate strategy is to leverage pre-existing molecular recognition of proteins for analyte specificity, yet attaching proteins to SWCNT for nanosensor generation remains challenging. Towards this end, we introduce a generalizable platform to generate protein-SWCNT-based optical sensors and use this strategy to synthesize a hydrogen peroxide (H_2_O_2_) nanosensor by covalently attaching horseradish peroxidase (HRP) to the SWCNT surface. We demonstrate a concentration-dependent response to H_2_O_2_, confirm the nanosensor can image H_2_O_2_ in real-time, and assess the nanosensor’s selectivity for H_2_O_2_ against a panel of biologically relevant analytes. Taken together, these results demonstrate successful covalent attachment of enzymes to SWCNTs while preserving both intrinsic SWCNT fluorescence and enzyme function. We anticipate this platform can be adapted to covalently attach other proteins of interest including other enzymes for sensing or antibodies for targeted imaging and cargo delivery.

## 1. Introduction

Single-walled carbon nanotubes (SWCNTs) possess unique optical and physical properties that make them attractive materials for biomedical applications.[1] In particular, their intrinsic photoluminescence in the near-infrared (nIR) region and lack of photobleaching are ideal traits for *in vivo* and *ex vivo* sensing and imaging in biological systems.[2,3] Another advantage of SWCNTs as nanosensors is the diversity of functionalization that add synergistic functions to the SWCNT conjugates. For instance, the functionalization of SWCNTs with biomolecules, such as single-stranded DNA (ssDNA), creates a corona phase at the proximity of SWCNT surfaces which can interact specifically with analytes of interest, resulting in modulation of the SWCNT optical signal.[4] Owing to their sequence modularity, various ssDNA sequences have been identified to play this corona phase molecular recognition role through screening for selective recognition of target molecules.[5] Furthermore, ssDNA can be evolved to have a selective interaction with an analyte through systematic evolution of ssDNA ligands by exponential enrichment.[6] While these ssDNA-SWCNT conjugates have shown promising analyte-specific fluorescence signal modulation, the platform fails to guarantee nanosensor generation for a specific analyte of interest as the molecular recognition element of these nanosensors is not rationally designed. On the other hand, the use of pre-existing molecular recognition elements such as proteins is an approach to rationally design selective SWCNT fluorescence modulation. Given their natural affinity for binding a target analyte, proteins are often the most readily available molecular recognition elements with which to develop nanosensors through protein-SWCNT conjugation.[7] While protein-SWCNT nanosensors can be rationally designed to detect or image a variety of biomarkers, the major drawback is that protein attachment to SWCNTs often compromises protein stability or SWCNT fluorescence.[8–10]

Traditionally, protein-SWCNT conjugates have been prepared by non-covalent approaches.[11] These methods typically rely on physical adsorption of the hydrophobic domains of proteins to SWCNT surfaces. For example, ultrasonication of SWCNT and protein[12] or dialysis-based ligand exchange[13] facilitates the non-specific adsorption of proteins to SWCNT surfaces. These non-covalent approaches, however, are dependent on the nature of the proteins which leads to varying levels of adsorption; thus, they are not generalizable.[14] Moreover, non-covalent attachment is often accompanied by conformational changes in the proteins, leading to the loss of their biological functions.[11] For example, Palwai *et al.* reported a complete loss of enzymatic activity of horseradish peroxidase (HRP) five days after immobilization on SWCNT.[15] A random alignment of the proteins on SWCNT may also reduce the efficiency of their biological activities, even if conformational stability is maintained. Therefore, non-covalent approaches are not a generalizable approach to developing protein-SWCNT nanosensors.

In this regard, covalent functionalization of proteins to SWCNT promises better stability and controlled alignment of the conjugated proteins.[16,17] However, covalent functionalization of SWCNT for sensing and imaging applications has been challenging because covalent bonds introduce sp^3^ defects into the sp^2^ SWCNT lattices. These sp^3^ quantum defects often attenuate or fully eliminate the optical transitions in SWCNTs that drive SWCNT photoluminescence.[18,19] On the other hand, recent reports have shown successful covalent conjugation of proteins and peptides to SWCNTs while preserving their optical properties. For example, it has been shown that minimal introduction of quantum defects can preserve or even enhance the photoluminescence of SWCNTs.[20,21] Following these findings, Mann *et al*. developed a protocol to covalently conjugate proteins to SWCNT by controlling the density of quantum defects.[22,23] An alternative approach is to covalently functionalize SWCNTs without introducing sp^3^ defects using azide-based conjugation.[24,25] Since the azide-based covalent bonds do not introduce an sp^3^ defect by re-aromatizing the sp^2^ lattice, this approach has a significantly higher degree of freedom in the density of functionalization compared to the approaches that use quantum defects. Recently, we have applied this chemistry to develop a versatile protocol to covalently functionalize SWCNTs and showed the potential of this chemistry to maintain analyte-specific responses of previously-reported nanosensors.[26]

In this study, we expand azide-based conjugation to develop new protein-based nanosensors by employing HRP as a model protein for the detection of hydrogen peroxide (H_2_O_2_). H_2_O_2_ is a critical component of reactive oxygen species (ROS) that play a pivotal role in many industrial and biological processes including ecosystem regulation in surface water, sterilization in food and beverage products, and cellular oxidative stress and signaling.[27,28] Consequently, developing fluorescent nanosensors for H_2_O_2_ that can be used for hydrogen peroxide detection or imaging is valuable, particularly non-photobleaching probes. To this end, we first developed and optimized a protocol to generate covalent HRP-SWCNT conjugates using azide-based triazine chemistry. We demonstrate that covalent conjugation does not compromise the photoluminescence properties of SWCNTs or the enzymatic activity of HRP, resulting in a robust turn-on fluorescence modulation upon nanosensor exposure to H_2_O_2_. Finally, we immobilized the nanosensors to show their potential applicability in bioimaging applications. Our results confirm that azide-based triazine chemistry can be used to develop new nanosensors via direct protein-SWCNT conjugation in a potentially generalizable platform, without compromising SWCNT fluorescence.

## 2. Results and Discussion

### 2.1. Nanosensor Platform Generation and Characterization

To synthesize our nanosensors, we first covalently modified unfunctionalized (pristine) SWCNTs with the azide-based triazine approach as previously reported (**Figure 1**a).[24,26] Briefly, we reacted pristine SWCNTs with cyanuric chloride and sodium azide to produce high density triazine-labelled SWCNTs (Trz-H-SWCNTs) with minimal quantum defects, maintaining intrinsic SWCNT optical properties like nIR fluorescence emission. Trz-H-SWCNTs were further functionalized with the amino acid cysteine in the presence of triethylamine. Nucleophilic substitution of the solvent-exposed chlorines on the triazine handles with the primary amine of cysteine produced thiol-functionalized SWCNTs (SH-SWCNTs). These SWCNTs maintained their optical properties as shown by the preservation of characteristic absorbance (Figure 1c) and fluorescence (Figure 1d) peaks. Notably, the choice of cysteine for functionalization was made in part due to Sulfo-SMCC being chosen as the crosslinker for this protein-SWCNT conjugation. This crosslinker is optimal for this conjugation scheme as it features two orthogonal functional groups: an N-hydroxysuccinimide-ester group (NHS-ester) that first reacts with solvent-exposed primary amines on HRP and a maleimide group that subsequently forms a stable covalent bond with free thiol groups on the SWCNT. This stepwise order of crosslinking afforded by Sulfo-SMCC helps minimize unwanted side reactions and imparts the flexibility of extending this platform to conjugate other proteins with exposed primary amines with the same SH-SWCNT sample.

**Figure 1.**
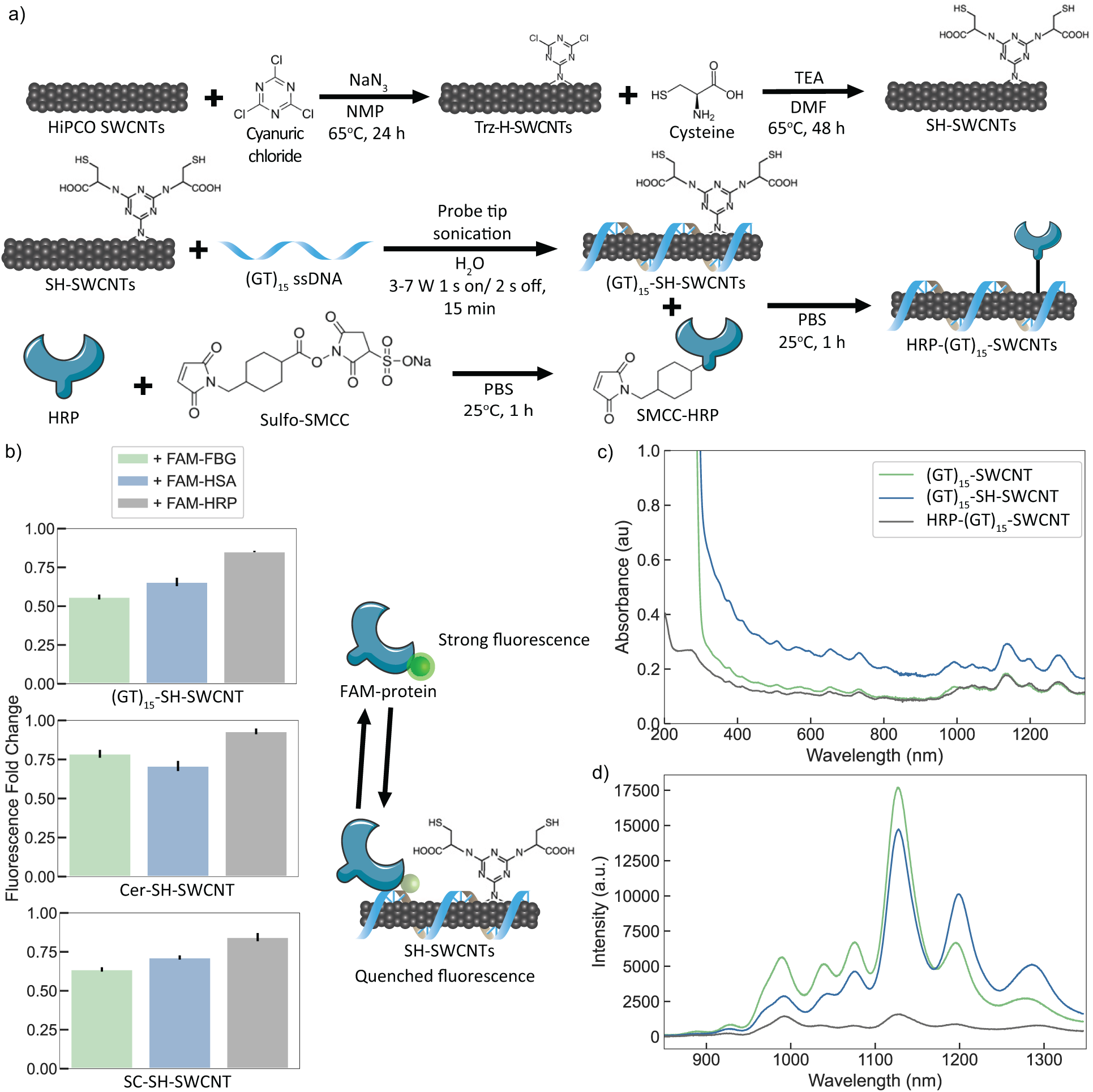
Synthesis and characterization of HRP-(GT)_15_-SWCNT nanosensors. (a) Synthesis scheme for nanosensor performed according to previously established protocols. The SMCC:HRP ratio was optimized for single maleimide addition (**Figure S1**) and reaction time and temperature were selected according to manufacturer protocol. Excess reactants were filtered away after each step and the final product was diluted back to the starting reaction volume. (b) Corona exchange assay for HRP and control proteins shows minimal HRP absorbance to different dispersant-SH-SWCNT samples. Fold change was calculated as the endpoint FAM fluorescence value 60 min after SWCNT addition to FAM-proteins divided by the initial FAM fluorescence for each protein sample. Error bars represent the standard deviation of experimental replicates (n = 3). (c) Absorbance spectra of (GT)_15_-coated SWCNTs along the nanosensor synthesis route show maintained characteristic peaks. (d) nIR fluorescence baselines for the samples in (c) show preservation of SWCNT intrinsic fluorescence throughout synthesis.

To solubilize SH-SWCNTs for conjugation with HRP, we assessed 3 potential amphiphilic SWCNT dispersants: (GT)_15_ ssDNA, the amphiphilic lipid C_16_-PEG2k-Ceramide (Cer), and the surfactant sodium cholate (SC). We induced noncovalent adsorption of each coating with SWCNTs through π-π aromatic stabilization and hydrophobic attraction, respectively. Each dispersant was added to SH-SWCNTs and subjected to probe-tip sonication on ice according to previously established protocols.[26,29] The resulting products ((GT)_15_-SH-SWCNT, Cer-SH-SWCNT, and SC-SH-SWCNT) showed high yield (100-300 mg/L) and solubility in water after centrifugation to remove aggregates and excess dispersant (**Figure S2**). The solubilized SWCNT products were subsequently assessed for the degree of nonspecific HRP adsorption to their surface with a corona exchange dynamics assay.[14] Briefly, this assay leverages the fluorescence quenching effect of fluorophores proximal to SWCNT to measure the degree of nonspecific protein adsorption to SWCNT surfaces. Compared to fluorescein (FAM)-functionalized control proteins fibrinogen (FBG-FAM) and human serum albumin (HSA-FAM), HRP-FAM showed lower FAM fluorescence quenching when incubated with SWCNTs, indicated by a higher endpoint fluorescence fold change value (Figure 1b). Since the degree of FAM quenching is proportional to the amount of nonspecific protein adsorption to the SWCNT surface, our results suggest that HRP shows minimal nonspecific adsorption to all three dispersed SWCNTs. This result highlights the utility of HRP as a model protein as its low level of adsorption helps ensure subsequent SWCNT sensor responses can be attributed to covalently attached HRP only, rather than a mixed population of covalently-attached and nonspecifically-adsorbed HRP. Though all three dispersants were good candidates in minimizing nonspecific HRP adsorption, we proceeded with H_2_O_2_ nanosensor development with (GT)_15_-SH-SWCNTs as they showed greater colloidal stability than SC-SH-SWCNTs through the rest of the sensor’s synthesis and greater response to H_2_O_2_ than Cer-SH-SWCNTs (**Figure S3**).

The covalent crosslinker Sulfo-SMCC was chosen to functionalize HRP with solvent-accessible maleimide groups for subsequent conjugation to (GT)_15_-SH-SWCNTs (Figure 1a). We reacted the NHS-ester group of Sulfo-SMCC with solvent-exposed primary amines on HRP to form maleimide-functionalized HRP (SMCC-HRP). After de-salting excess unreacted crosslinker, we reacted SMCC-HRP with (GT)_15_-SH-SWCNTs to covalently link HRP to SWCNTs. Following centrifugal membrane filtration to remove excess unreacted SMCC-HRP, we characterized the final nanosensor product (HRP-(GT)_15_-SWCNT). The nanosensor showed characteristic optical absorbance (Figure 1c) and fluorescence (Figure 1d) properties of SWCNT, with minimal changes to the SWCNT fluorescence peak wavelengths for HRP-(GT)_15_-SWCNT relative to the base (GT)_15_-SH-SWCNTs material. We did find that attaching HRP to SWCNTs yielded a decrease in fluorescence intensity, a desired and common mechanism to generate turn-on fluorescent nanosensors.[30] We hypothesize that the local dielectric environment of HRP-(GT)_15_-SWCNT is affected by the proximity of HRP to the SWCNT surface upon conjugation, leading to attenuated fluorescence intensity. If this attenuation was due to quantum defects or other sp^2^ lattice damage introduced by the synthesis process, the SWCNT fluorescence intensity would not be modulated in the presence of the HRP substrate H_2_O_2_ but instead remain attenuated. Subsequent experiments show that addition of H_2_O_2_ induces a return to previous baseline SWCNT intensities, pointing towards HRP as the source of the attenuated sensor baseline fluorescence. Additionally, previous studies have shown that SWCNT fluorescence intensity decreases in the presence of H_2_O_2_ rather than increase, providing further evidence that the catalysis of H_2_O_2_ by HRP facilitates the turn-on fluorescence response.[31]

After developing and optimizing the reaction conditions for this sensor, we characterized its physical and chemical properties to confirm the successful covalent attachment of HRP to SWCNTs while maintaining enzymatic activity. To visualize HRP on the surface of SWCNTs, we captured atomic force microscopy (AFM) images of HRP (**Figure 2**a) and HRP-(GT)_15_-SWCNT (Figure 2b). Height trace analysis along the length of the nanotube showed several peaks of increased height ∼0.9 nm greater than the height of the nanotube as determined by tracing perpendicular to the SWCNT length axis (Figure 2c). Combined with the corona exchange results confirming minimal nonspecific adsorption of HRP to SWCNTs, these peaks can thus be attributed to covalently attached HRP as shown schematically (Figure 2c).

**Figure 2.**
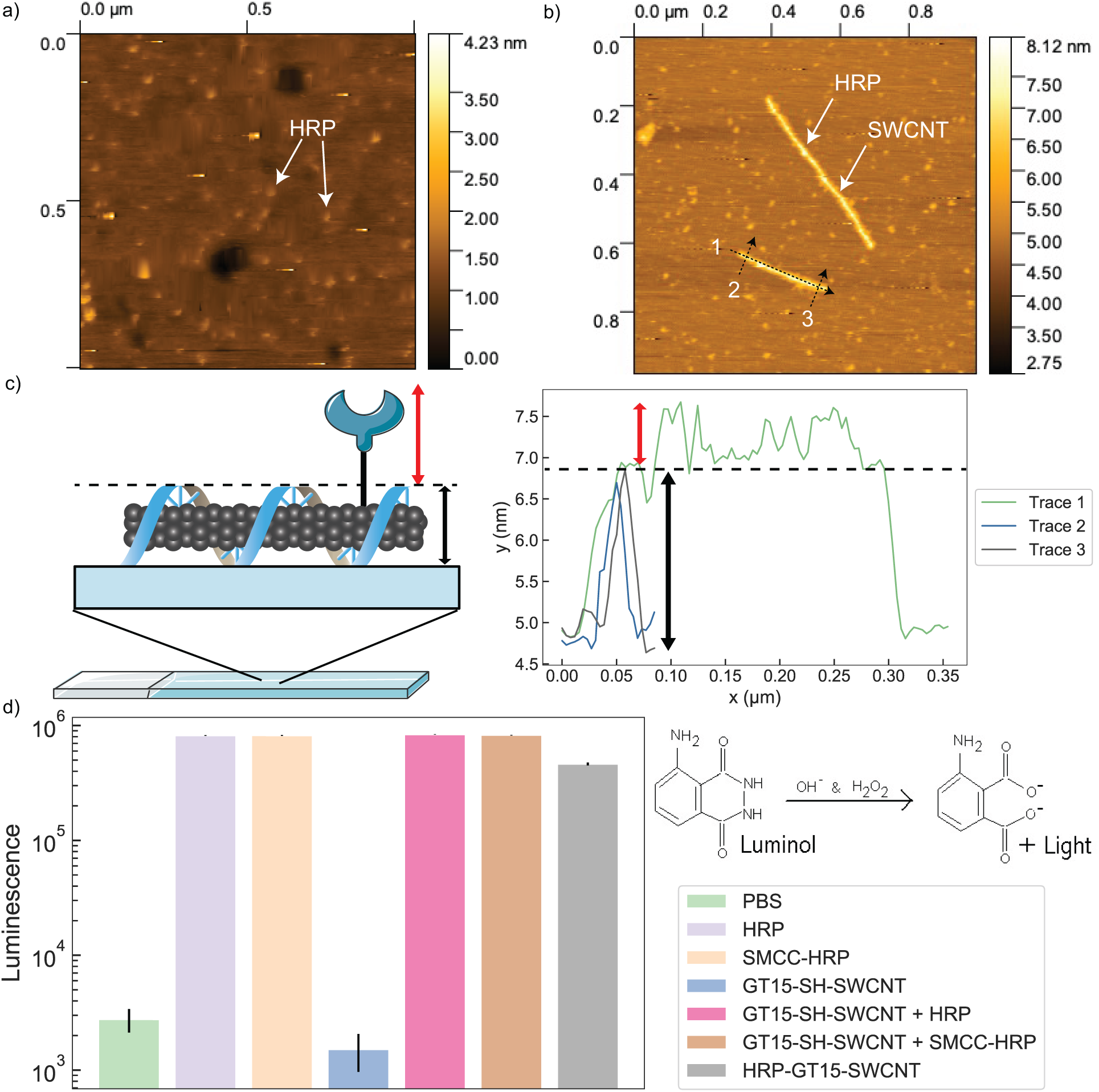
Validation of HRP presence and activity for HRP-(GT)_15_-SWCNT nanosensors. (a) AFM image of HRP showing a height increase of ∼0.9 nm for this protein. (b) AFM image of nanosensor shows individual SWCNTs with several height spikes along axis, visually confirming both successful (GT)_15_ dispersion (minimal aggregates) and HRP conjugation. (c) Height trace analysis of the AFM image in (b) shows that the height spikes along the SWCNT axis (Trace 1) correspond to HRP height above SWCNT baseline (Trace 2,3). (d) Luminol activity assay shows nanosensor has comparable HRP activity to control and that neither SMCC functionalization nor SWCNT presence interferes with HRP activity. Error bars represent the standard deviation between experimental replicates (n = 3).

To confirm that covalently attached HRP maintained enzymatic activity while on the surface of SWCNTs, we used a luminol assay according to established protocols.[32] Briefly, the oxidation of luminol by the catalysis of H_2_O_2_ by HRP produces luminescence proportional to protein activity when normalized for protein concentration. We thus measured luminescence values for all HRP conditions along the nanosensor synthesis route incubated with luminol and H_2_O_2_ (Figure 2d). Compared to the low luminescence magnitude shown by the control buffer and (GT)_15_-SH-SWCNT alone, the nanosensor showed similar catalytic activity to the other HRP-containing samples at the same protein concentration. This data also suggests that SMCC functionalization does not negatively affect HRP activity under these conditions. Similarly, the presence of SWCNT showed little negative impact on the activity of both native and SMCC-HRP when mixed. Taken together, these results suggest that we have synthesized a covalent HRP-SWCNT sensor with preserved SWCNT and enzymatic properties that can be assessed for its ability to sense hydrogen peroxide.

### 2.2. Characterizing Nanosensor Response to Analyte Hydrogen Peroxide

We characterized our HRP-(GT)_15_-SWCNT nanosensor’s response to hydrogen peroxide by measuring the change in nIR SWCNT fluorescence over time in response to varied levels of H_2_O_2_ analyte, schematically represented in **Figure 3**a. The addition of H_2_O_2_ elicited a strong and stable turn-on fluorescence response over the course of one hour (Figure 3c). This contrasts with the minor turn-off response elicited by water (Figure 3b), isolating the analyte as the cause of the turn-on response rather than the addition of volume to the sample. The normalized integrated change in fluorescence (ΔF/F_0_) of the sensor peaked after 20 min post-addition of H_2_O_2_ and remained stable over the course of one hour (Figure 3d). In contrast, (GT)_15_-SH-SWCNT alone exhibits a turn-off fluorescence response to H_2_O_2_ (**Figure S4**), as expected from previous literature.[31] Furthermore, HRP mixed with (GT)_15_-SH-SWCNT exhibits a strong turn-off response immediately upon H_2_O_2_ addition followed by a gradual increase in fluorescence to the baseline intensity (Figure S4). This could be due to the consumption of H_2_O_2_ by free HRP, mitigating the analyte’s quenching effect as it is depleted from the solution. These results suggest that the strong turn-on response of our sensor must be due to the interaction between H_2_O_2_ and covalently-attached HRP on the SWCNT surface rather than between H_2_O_2_ or catalysis reaction byproducts and the SWCNT itself.

**Figure 3.**
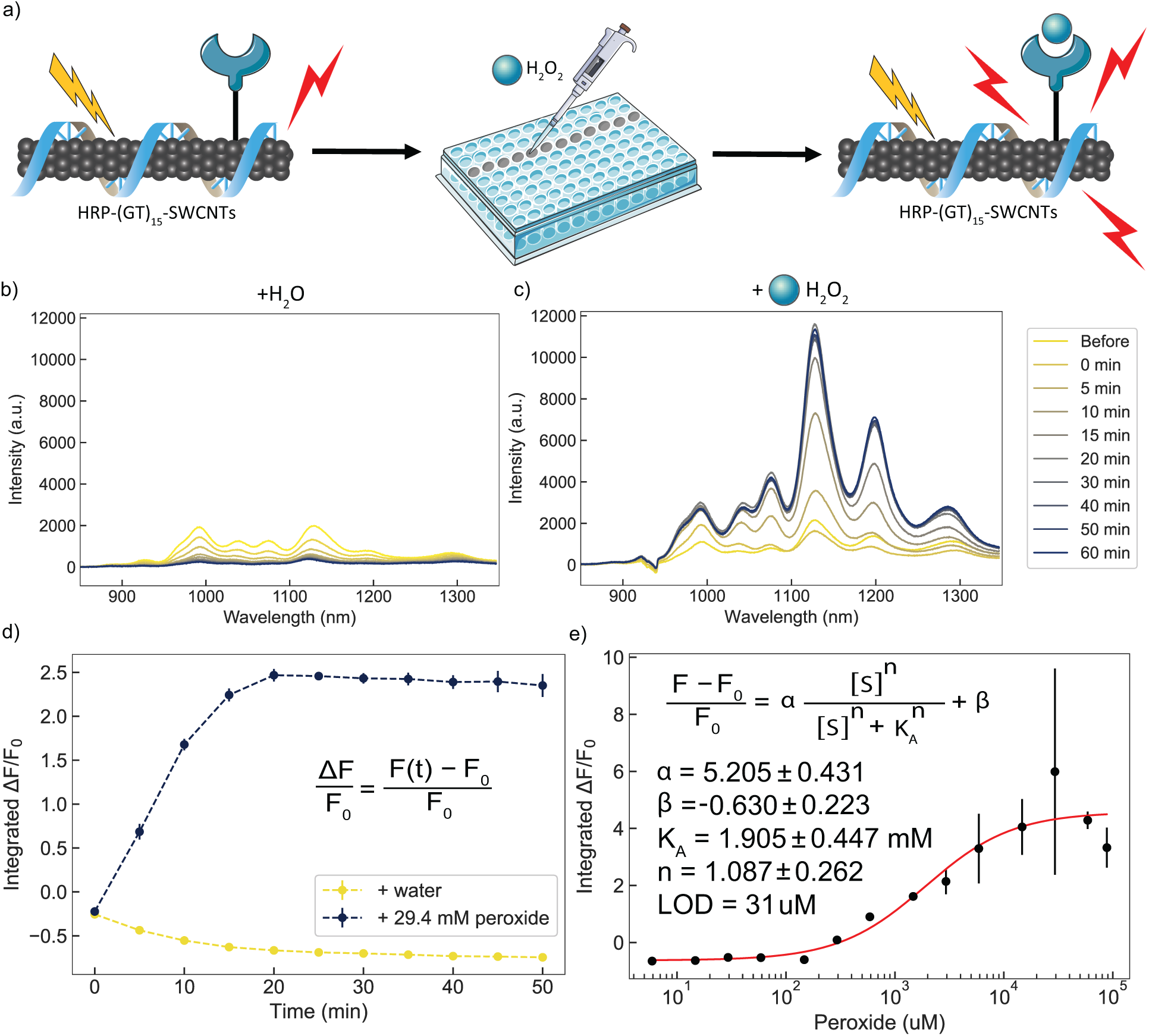
HRP-(GT)_15_-SWCNT nanosensor response to H_2_O_2_ in solution-phase screen. (a) Schematic depiction of response screening platform, where addition of analyte to a 384-well plate with nanosensor induces a strong turn-on response as measured by the increase in nIR fluorescence emission. (b) Nanosensor response to water shows a strong turn-off response over 60 min. (c) Nanosensor response to 29.4 mM H_2_O_2_ shows a strong turn-on response over 60 min. (d) The normalized integrated change in fluorescence (ΔF/F_0_) for (b) and (c). (e) Endpoint (time = 60 min) ΔF/F_0_ values for varying concentrations of analyte plotted and fit to a cooperative binding model. Fit parameters are listed with 95% confidence intervals evaluated using the t-distribution. All fluorescence measurements were obtained with a laser excitation wavelength of 721 nm. Error bars represent the standard deviation between experimental replicates (n = 3).

We further measured the concentration-dependent nanosensor response and determined a 31 μM limit of detection (LOD) for hydrogen peroxide (Figure 3e). The data was fit to a cooperative binding model and extracted parameters include an equilibrium constant K_A_ of 1.905 mM and Hill coefficient n of 1.087, as expected of this noncooperative enzyme which uses its heme cofactor to bind one peroxide molecule at a time for catalysis.[33] Since HRP bound to SWCNT shows the expected noncooperativity value of n=1, we can interpret the equilibrium constant K_A_ as the dissociation constant K_d_ interchangeably.[34]

Comparing these extracted parameters to previous literature, we observe a decreased K_d_ value and comparable LOD. Previous literature found free HRP to have a K_d_ of 3.4-4.4 mM, which is higher than our nanosensor’s value.[35,36] Excitingly, this result indicates that HRP covalently bound to SWCNTs shows higher affinity for H_2_O_2_ than free enzyme. This principle has been demonstrated in previous studies immobilizing enzymes to electrodes in electrochemical biosensors.[37–40] When compared with similar peroxide sensors, our nanosensor LOD was within the range of literature values (**Table 1**). Though our nanosensor has a higher LOD and linear range than some systems (**Figure S5**), our system maintains the advantage of facile synthesis, form factor variability, non-photobleaching nature, and an underlying platform that is flexible enough to accommodate other proteins and analytes to generate other nanosensors or multiplexed sensors.

**Table 1.** Comparisons of similar H_2_O_2_ nanosensors found in literature.

| Platform | Signal Type | Linear Range (μM) | LOD (μM) | Reference |
| --- | --- | --- | --- | --- |
| HRP/Os Polymer | Amperometric | 1-500 | 0.3 | [40] |
| HRP/Au | Amperometric | 40-100 | 40 | [41] |
| SWCNT/GCE | Amperometric | 1900-24000 | 1000 | [42] |
| HRP/Luminol | Chemiluminescence | 100-3000 | 667 | [43] |
| Fe-N-C Nanzyme | Chemiluminescence | 500-100000 | 0.5 | [44] |
| Cobalt-CNT/GOx | Fluorescence | 0.2-20 | 0.1 | [45] |
| ssDNA-SWCNT | Fluorescence | 10-1000 | 0.1 | [31] |
| AgNP | Absorbance | 10-10000 | 10 | [46] |
| HRP-SWCNT | Fluorescence | 150-600 | 31 | This work |

Abbreviations: Ag, silver; Au, gold; Fe, iron; N, nitrogen; C, carbon; GCE, glassy carbon electrode; GOx, glucose oxidase, Os, Osmium.

### 2.3. Immobilized Nanosensor Response to Analyte

After characterizing the solution-phase nanosensor response, we investigated the nanosensor response when surface-immobilized on a glass microwell dish (**Figure 4**a). HRP-(GT)_15_-SWCNTs were immobilized on a glass microwell dish and imaged after rehydration with PBS over the course of 5 min. Isolating the image analysis to fluorescent HRP-(GT)_15_-SWCNT regions of interest with high fluorescence and averaging their values over time, we demonstrate the nanosensor’s ability to sense hydrogen peroxide when immobilized through an imaging format (Figure 4b). The addition of water at 60 s shows no measurable fluorescence change, while the addition of 588 μM H_2_O_2_ at 120 s shows a signal increase of ∼15%, with representative images at Frame 9 (Figure 4c) and 124 (Figure 4d). This response can thus still be attributed to the analyte’s presence upon addition rather than volume increases. Further, the repeatability of this response can be seen by the similar sharp increases in fluorescence upon subsequent additions of hydrogen peroxide every 120 s (Figure 4e). Though the signal fails to fully return to baseline between analyte additions likely due to imaging drift, the similar increase in magnitude for each peak suggests that the consumption of analyte by HRP is modulating SWCNT fluorescence intensity rather than interaction between the analyte or reaction byproducts and the SWCNT itself. The difference in magnitude of fluorescence modulation between the solution phase HRP-(GT)_15_-SWCNT nanosensor response to hydrogen peroxide and this immobilized form factor can be attributed to the difference in analysis between the two. In solution, the fluorescence is measured for each wavelength in the nIR region, whereas the immobilized form factor captures fluorescence images of the sample, aggregating the total emission over the nIR region into an image. The solution-phase response is stronger at certain nIR wavelengths (Figure 3c), corresponding to different chirality SWCNTs. This sensitivity is lost in the images as they only capture overall emission, limiting the magnitude of the response to the same concentration of H_2_O_2_.

**Figure 4.**
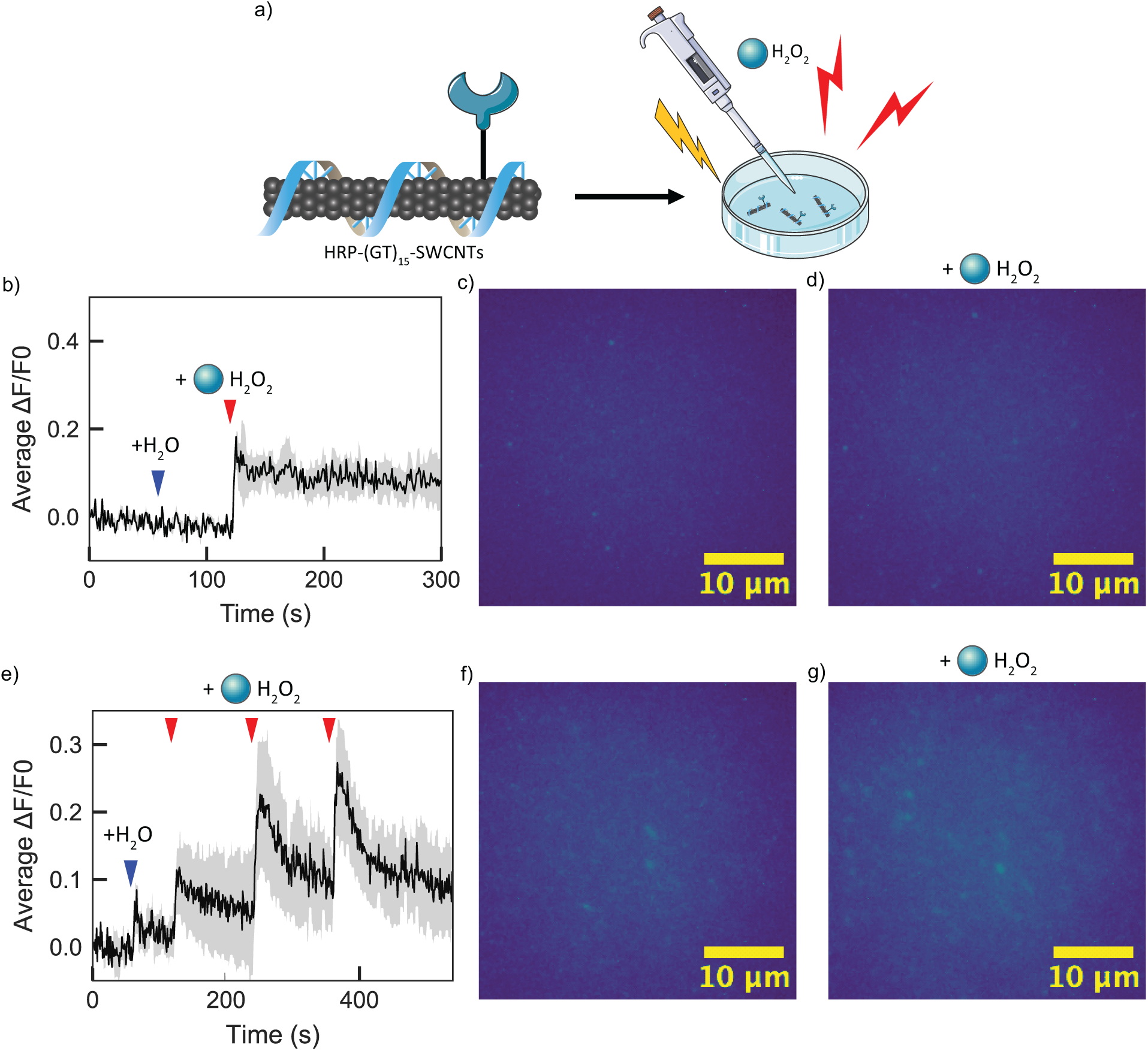
Immobilized nanosensor response to H_2_O_2_. (a) Schematic depiction of immobilized sensing platform, where the nanosensor is incubated in a glass microwell dish for 20 min, washed, and rehydrated with 1X PBS before analyte addition and imaging. Addition of analyte induces an increase in fluorescence intensity of the sensor. (b) Analysis of fluorescence microscopy images of immobilized nanosensor over 5 min shows around 15% increase in fluorescence intensity corresponding to H_2_O_2_ addition at 120 s. (c) Fluorescence microscopy image of immobilized nanosensor at 9 s post-hydration with PBS. (d) Fluorescence microscopy image of immobilized nanosensor at 4 s post-addition of H_2_O_2_ shows increased fluorescence intensity compared to (c). (e) Analysis of fluorescence microscopy images of immobilized nanosensor over 10 min shows repeatable fluorescence intensity increase upon subsequent additions of H_2_O_2_. (f) Fluorescence microscopy image of immobilized nanosensor at 9 s post-hydration with PBS. (g) Fluorescence microscopy image of immobilized nanosensor at 4 s post the third addition of H_2_O_2_ shows increased fluorescence intensity compared to (f). Gray shaded areas represent the standard deviation from the mean value in black (n=20). Average fluorescence change values in (b) and (e) are calculated using ROIs from the entire image field of view. Images in (c), (d), (f), and (g) are representative images highlighting the center of the field of view.

### 2.4. Nanosensor Analyte Selectivity

Following successful demonstration of the surface-immobilized nanosensor, we next characterized the HRP-(GT)_15_-SWCNT nanosensor’s selectivity for H_2_O_2_ against a panel of relevant analytes (**Figure 5**a). These analytes were chosen to both ensure that the catalysis of H_2_O_2_ was the sole signal source rather than similar structural analogues (TBHP) or downstream reaction intermediates (superoxide), and to evaluate the potential interference of biomolecules that would be present during *in vitro* sensing applications (glutathione and sodium hypochlorite). Compared to buffer and H_2_O_2_ control, the addition of 50 μM of GSH, NaOCl, and TBHP produced either no response or a turn-off response. The addition of 50 μM of superoxide did produce a turn-on response similar to H_2_O_2_, though this result is expected as superoxide bound to the heme core of HRP is an intermediate of the enzyme’s catalytic cycle (Compound III).[47,48] Altogether, our nanosensor appears to be selective for ROS over other potential interfering species, further supporting the necessity of HRP-analyte interaction for the observed fluorescence turn-on response. Further relevant analytes can be quickly assayed using the immobilized form factor depending on the intended application.

**Figure 5.**
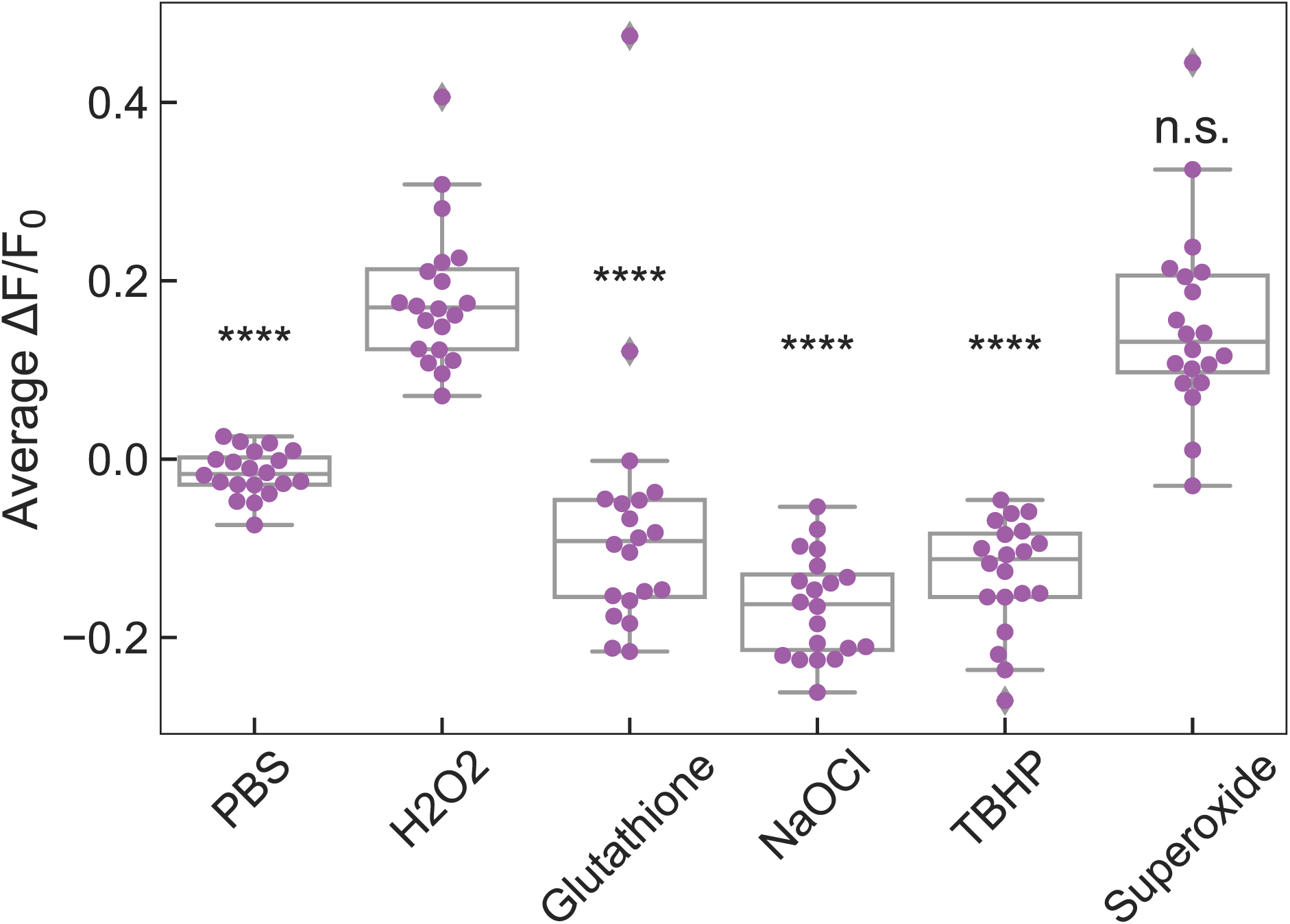
Nanosensor selectivity against relevant analytes. Normalized change in fluorescence values of nanosensor immobilized on glass upon addition of analytes shows minimal response for analytes other than the target. Fluorescence change for each ROI (n=20) 10 s after analyte addition is plotted as purple circles for each condition. ****P = 3.35 x 10^-9^ (PBS), 1.64 x 10^-13^ (Glutathione), 2.26 x 10^-14^ (NaOCl), and 2.26 x 10^-14^ (TBHP), and P = 0.89 (Superoxide) in independent two-sample *t* tests for each analyte ΔF/F_0_ in comparison to H_2_O_2_.

## 3. Conclusion

The generation of SWCNT-based nanosensors has been important to enable the visualization of biological analytes through the unique near-infrared fluorescence of SWCNTs. One outstanding challenge in the development of SWCNT-based nanosensors lies in identifying suitable molecular recognition elements to provide analyte-selective modulation of SWCNT fluorescence. Direct attachment of protein-based molecular recognition agents would enable a design-based approach to nanosensor development so long as protein function and SWCNT intrinsic fluorescence can both be preserved. In this work, we show that triazine-based SWCNT functionalization can be used to attach HRP enzymes to SWCNTs and demonstrate H_2_O_2_ sensing as a proof-of-principle demonstration of nanosensor functionality. To generate these sensors, we leveraged triazine-based SWCNT functionalization to attach free thiol groups to pristine SWCNTs while maintaining their optical properties. We determined that the resulting HRP-(GT)_15_-SWCNT nanosensor showed a concentration-dependent turn-on fluorescence response to H_2_O_2_ in solution, partly due to the quenched SWCNT fluorescence baseline intensity after HRP conjugation. Fitting this response to a cooperative binding model produced estimated kinetic parameters including a solution-phase LOD of 31 μM. Though this value is above typical cellular ROS levels (100 nM-10 μM),[49] the nanosensor showed the ability to sense H_2_O_2_ when surface-immobilized on glass, presenting an alternate viable sensing form factor for effective H_2_O_2_ sensing in other systems with higher H_2_O_2_ levels such as contaminated water samples and food production.[28] In this form factor, the nanosensor showed robust analyte selectivity against similar analytes. Taken together, this study demonstrates the potential of covalent protein-SWCNT nanosensors for sensitive, specific, and stable analyte sensing.

The nanosensor synthesis platform developed here can also be extended to conjugate other proteins and enzymes of interest and generate robust nanosensors for biologically relevant analytes. The use of Sulfo-SMCC as the crosslinking agent allows any protein with exposed primary amines to be a candidate for conjugation to SH-SWCNTs. Additionally, the use of Triazine-SWCNT chemistry provides a wide library of available SWCNT functional groups like carboxylic acid (-COOH), primary amine (-NH_2_), and biotin. These functionalized SWCNTs could thus be used with other crosslinker systems to conjugate proteins that are not amenable to maleimide functionalization via Sulfo-SMCC. However, future work in this area would require reaction condition optimization to ensure product sensors are colloidally stable and enzymatically active. Orthogonally, SWCNTs are also confirmed to be valuable as biomolecule delivery agents,[50–53] and our conjugation work herein could enable protein conjugation to SWCNTs for protein delivery applications. Ultimately, these nanosensors could help advance the use of SWCNTs as nanomaterials for theranostics, delivery, and cellular fluorescence imaging purposes.

## 4. Experimental Section

### Materials

All chemicals unless otherwise stated were purchased from Sigma-Aldrich. Raw high pressure carbon monoxide (HiPco) synthesized SWCNTs were purchased from NanoIntegris. C_16_-PEG2k-Ceramide (N-palmitoyl-sphingosine-1-{succinyl[methoxy(polyethyleneglycol)2000]}) phospholipid was purchased from Avanti Polar Lipids. (GT)_15_ ssDNA was purchased from Integrated DNA Technologies. Sulfo-SMCC (sulfosuccinimidyl 4-(N-maleimidomethyl)cyclohexane-1-carboxylate) was purchased from ThermoFisher Scientific. Luminol (Pierce ECL Western Blotting Substrate) was purchased from ThermoFisher Scientific. Glass-bottom microwell dishes (35 mm petri dish with 10 mm microwell) were purchased from MatTek. Hydrogen Peroxide (3% w/w) was purchased from Thomas Scientific. Sodium hypochlorite was purchased from Thomas Scientific. Tert-butyl hydroperoxide (TBHP) was purchased from Thomas Scientific. Xanthine was purchased from Thomas Scientific. Xanthine Oxidase (from buttermilk) was purchased from Millipore Sigma. 2-(2-methoxy-4-nitrophenyl)-3-(4-nitrophenyl)-5-(2,4-disulfophenyl)-2H-tetrazolium (WST-8) was purchased from ThermoFisher Scientific.

### Synthesis of SH-Functionalized SWCNTs

Synthesis of SH-functionalized SWCNTs was adapted from previous literature.[24,26] Briefly, pristine HiPco SWCNTs (1 g) were dispersed in *N*-methyl-2-pyrrolidone (NMP) (150 mL) in a round bottom flask with a stir bar. The mixture was bath sonicated (Branson Ultrasonic 1800) for 1 h at 25 °C followed by gentle stirring for 1 h at 25 °C. The mixture was then cooled to 0 °C on ice. 2,4,6-1,3,5-trichloro-triazine (10 g, 54 mmol) was dissolved in NMP (50 mL) and slowly added to the SWCNT dispersion. The resulting mixture was stirred for 20 min at 0 °C. Sodium azide (1.76 g, 27 mmol) was gradually added to the mixture and stirred for 2 h at 0 °C. The mixture was then stirred at 25 °C for 1 h followed by stirring at 70 °C for 12 h to yield Trz-H-SWCNTs. This product was purified by repeated centrifugation and washing steps with 10 mL each of acetone, water, toluene, then chloroform. The purified product was then lyophilized for storage and characterization.

Trz-H-SWCNTs (10 mg) were dispersed in dimethylformamide (DMF) (5 mL) and bath sonicated for 15 min at 25 °C. Cysteine (1 mg) and a 1.5 M excess of triethylamine were added to the mixture and stirred for 48 h at 65 °C. The product (SH-SWCNTs) was purified by centrifugation, supernatant removal, and re-dispersion in washes of DMF (4 mL, 2X) then water (4 mL, 2X). The product was then dialyzed against water using a Slide-A-Lyzer G2 10 kDa MWCO dialysis cassette (Thermo Scientific) for 1 week with daily water changes (2 L). The purified product was pelleted by centrifugation and lyophilized for storage, characterization, and covalent HRP functionalization.

### Noncovalent Adsorption of ssDNA to SH-SWCNTs by Probe-Tip Sonication

SH-SWCNTs (1 mg) and (GT)_15_ single-stranded DNA (1 mg) were dispersed in 1X phosphate buffered saline (PBS) (500 μL, pH 7.4) and bath sonicated for 10 min at 25 °C. The solution was then probe-tip sonicated with an ultrasonic processor (Cole Parmer) and a 3 mm stepped microtip probe with pulses of 3-7 W for 1 s followed by 2 s of rest for a total sonication time of 15 min. The solution was equilibrated for 1 h at 25 °C then subsequently centrifuged at 16100 relative centrifugal force (RCF) for 30 min to remove unsuspended SWCNT aggregates. Suspended SWCNTs formed a homogeneous dark gray solution and were measured for concentration by UV-Vis-IR absorbance (Shimadzu UV-3600 Plus) with samples in a 100 μL volume, black-sided quartz cuvette (Thorlabs, Inc.). SWCNT concentration was calculated from absorbance at 632 nm using the Beer-Lambert law with extinction coefficient ε_632_ = 0.036 L mg^-1^ cm^-1^.[54]

### Corona Exchange Dynamics Assay for HRP-SWCNT Adsorption

Corona exchange dynamics studies were conducted as described previously.[14] Briefly, HRP, fibrinogen, and human serum albumin were labeled with a fluorophore (FAM) via N-hydroxysuccinimide ester conjugation (Lumiprobe). Protein (10 mg) in 1X PBS (900 μL) and an 8-fold molar excess of FAM-NHS in dimethyl sulfoxide (DMSO) (100 μL) were gently mixed via end-over-end rotation in a foil-covered tube for 4 h. FAM-protein conjugates were then purified with Zeba 2 mL spin desalting columns with 40 kDa MWCO (Thermo Scientific) to remove excess unreacted FAM-NHS according to manufacturer’s instructions. The purified FAM-proteins were measured for concentration and degree of labeling via UV-Vis-IR absorbance at 280 nm for protein and 495 nm for FAM. The degree of labeling was calculated as the molar ratio of FAM to protein in the samples. 200 mg L^-1^ FAM fluorophore-labeled protein (25 μL) was added to 10 mg L^-1^ SH-SWCNTs dispersed with (GT)_15_ ssDNA, C_16_-PEG2k-Ceramide, and SC (25 μL) in triplicate. The solutions were combined via microchannel pipette in a 96-well PCR plate (Bio-Rad) and mixed by pipetting. The plate was sealed with an optically transparent adhesive seal and gently spun down in a benchtop centrifuge. Fluorescence time series measurements were obtained with a Bio-Rad CFX96 Real Time qPCR System by scanning the FAM channel every 30 s at 25 °C.

### Covalent Conjugation of HRP to SH-SWCNT via Sulfo-SMCC Crosslinker

(GT)_15_-SH-SWCNTs were diluted to 20 mg L^-1^ in PBS and 5% v/v TCEP to reduce disulfide bonds between SWCNTs. HRP was similarly diluted to 2 mg mL^-1^ in PBS and Sulfo-SMCC dissolved in Milli-Q H_2_O was added at 10:1 molar ratio Sulfo-SMCC:HRP. Both mixtures were incubated separately for 1 h at 25 °C. Each solution was then de-salted to remove excess TCEP and Sulfo-SMCC, respectively, with 7K MWCO Zeba Spin Desalting Columns (2 mL) according to manufacturer’s instructions. Purified (GT)_15_-SH-SWCNTs and maleimide-functionalized HRP were then mixed and incubated 1 h at 25 °C. The finished reaction mixture was purified to remove unconjugated maleimide-HRP with 100K MWCO Amicon Ultra Centrifugal Filters (0.5 mL). The membrane was rinsed with PBS and centrifuged at 5000 rcf for 5 min. The raw reaction mixture was then added and spun at the same conditions. PBS (450 μL) was added to the membrane to wash away excess protein and spun at the same conditions and repeated once. Finally, the membrane was cleaned with PBS (200 μL) to remove adsorbed SWCNTs and the membrane was inverted and spun at 1000 rcf for 2 min to collect the purified sample. The recovered HRP-(GT)_15_-SWCNTs were then diluted to their original reaction volume and characterized by UV-Vis-IR absorbance and nIR fluorescence measurements.

### Atomic Force Microscopy of SWCNT Nanosensors

SWCNT nanosensors were analyzed for the presence of HRP with atomic force microscopy (AFM). A small square (1 cm x 1 cm) of mica substrate was adhered to a glass slide and the top surface was freshly cleaved with tape immediately prior to sample preparation. HRP-(GT)_15_-SWCNT (100 μL) at 5 mg L^-1^ in 1X PBS was statically dispensed onto the mica surface and spin-coated at 2000 rpm for 1 min. Static dispense and spin coating was then repeated with an additional 100 μL of 5 mg L^-1^ HRP-(GT)_15_-SWCNT to increase the coverage of functionalized SWCNT on the mica surface. Once doubly coated, deionized water (100 μL) was slowly dynamically dispensed onto the coated surface while spinning to rinse off excess salts. The sample was then stored at room temperature overnight and imaged using a TAP150AL-G (Ted Pella) Aluminum Reflex coated tip coupled to an MFP-3D-BIO AFM (Asylum Research) in soft tapping mode.

### Luminol Assay for HRP Activity

HRP activity was assessed using the Pierce^TM^ ECL Western Blotting Substrate kit. Briefly, the samples were all diluted to the same HRP concentration as the lowest concentration to be measured for activity, typically the nanosensor. Each sample was confirmed to be at the correct [HRP] via Qubit assay according to manufacturer’s instructions. The peroxide and luminol stocks from the kit were mixed at 1:1 by volume and added to each sample to dilute the [HRP] to the working concentration of 0.5 mg L^-1^ according to the manufacturer’s instructions. Each sample (50 μL) was added to a 96-well plate in triplicate and the luminescence of each well was measured over 60 min on a luminescence plate reader (Tecan M1000).

### Near-Infrared Spectroscopy of SWCNT Nanosensors

Fluorescence spectra were obtained with an in-house nIR microscope setup. Briefly, we used an inverted Zeiss microscope (Axio Observer.D1, 10X objective) coupled to a Princeton Instruments spectrometer (SCT 320) and liquid nitrogen-cooled Princeton Instruments InGaAs detector (PyLoN-IR). SWCNT samples were excited with a 721 nm laser (OptoEngine LLC) and emission was collected in the 850 – 1350 nm wavelength range. The samples were measured in a 384 well-plate (Greiner Bio-One microplate) with a total volume of 30 μL per well. For solution-phase sensor response screens, nanosensor at 2.5 mg L^-1^ [SWCNT] in 1X PBS (27 μL) was added per well and 10X H_2_O_2_ (3 μL) was injected per well with a microchannel pipette in triplicate. After analyte addition, each well was briefly mixed by pipetting, sealed with an adhesive seal (Bio-Rad) and spun down for 10 s with a benchtop well plate centrifuge to remove bubbles. Fluorescence spectra were measured at time points of 0 min after analyte addition, 5 min, 10 min, 15 min, 20 min, and every 10 min after until 1 h post-addition.

### Near-Infrared Microscopy of SWCNT Nanosensors

Fluorescence images were captured with the same epifluorescence microscope setup as described previously with a 100X oil immersion objective and a Ninox VIS-SWIR 640 camera (Raptor). Nanosensors were immobilized on glass-bottom microwell dishes (35 mm petri dish with 10 mm microwell, MatTek) as follows: each dish was washed twice with PBS (150 μL), incubated with nanosensor at 2.5 mg L^-1^ (100 μL) for 20 min, and washed twice again with PBS (150 μL). For each image, PBS (160 μL) was added to the dish and the z-plane was refocused to maximize SWCNT fluorescence intensity. Images were then recorded over 5 min with an exposure time of 950 ms and 1000 ms repeat cycle. Water (20 μL) was added at Frame 60 and analyte (20 μL) was added at Frame 120. Images were processed with ImageJ as follows: a median filter (0.5-pixel radius) and rolling ball background subtraction (300-pixel radius) were applied, the image was cropped to eliminate gaussian blur and highlight the center of the image with the brightest nanosensors (width = 375, height = 375, x-coordinate = 110, y-coordinate = 36), the image was then analyzed using the ROI analyzer tool (Multi Measure) highlighting the clearest 20 ROIs of nanosensor bundles.

### Superoxide Generation

Superoxide was generated enzymatically with the Xanthine/Xanthine Oxidase system according to previously established protocols.[48] Briefly, Xanthine was dissolved at 10 mM in 0.1 M NaOH and the pH was adjusted to 7 with 0.1 M HCl and a pH probe. Xanthine Oxidase was diluted to 0.1 U mL^-1^ in 1X PBS. Xanthine and Xanthine Oxidase (50 μL each) were added to 1X PBS (140 μL) in an Eppendorf tube and incubated 2 h at 25 °C. Separately, the generation of superoxide was validated by incubating 20 mg mL^-1^ WST-8 with the reaction mixture. The absorbance at 460 nm of the resulting solution was measured to determine the presence of the WST-8 formazan product, which is proportional to the generation of superoxide. Using the Beer-Lambert law with extinction coefficient ε_460_ = 30,700 M^-1^ cm^-1^ for the formazan product and knowing 2 superoxide radicals are required to generate 1 formazan, we calculated the superoxide concentration in this system (**Figure S6**).[55]

## Supporting Information

Supporting information is available from the Wiley Online Library or from the author.

## Acknowledgements

M.P.L. acknowledges the support of Burroughs Wellcome Fund Career Award at the Scientific Interface (CASI), the Simons Foundation, NIH NIDA CEBRA award #R21DA044010, Stanley Fahn PDF Junior Faculty Grant with Award #PF-JFA-1760, Beckman Foundation Young Investigator Award, DARPA Young Faculty Award, FFAR New Innovator Award, an IGI award, support from CITRIS and the Banatao Institute, and a USDA award. M.P.L. is a Chan Zuckerberg Biohub Investigator and an Innovative Genomics Institute Investigator. F.L. acknowledges the support of the NSF Graduate Research Fellowship (NSF DGE 1752814). The authors would like to thank Rebecca L. Pinals, Jaquesta A.M. Adams, Natsumi Komatsu, Jeffrey W. Wang, and Jaewan Mun for insightful conversation and feedback. The authors would like to acknowledge the use of clipart from Servier (smart.servier.com).

## Notes

### Competing Interest Statement

The authors have declared no competing interest.

