## Supporting Information for "Covalent Attachment of Horseradish Peroxidase to Single-Walled Carbon Nanotubes for Hydrogen Peroxide Detection"

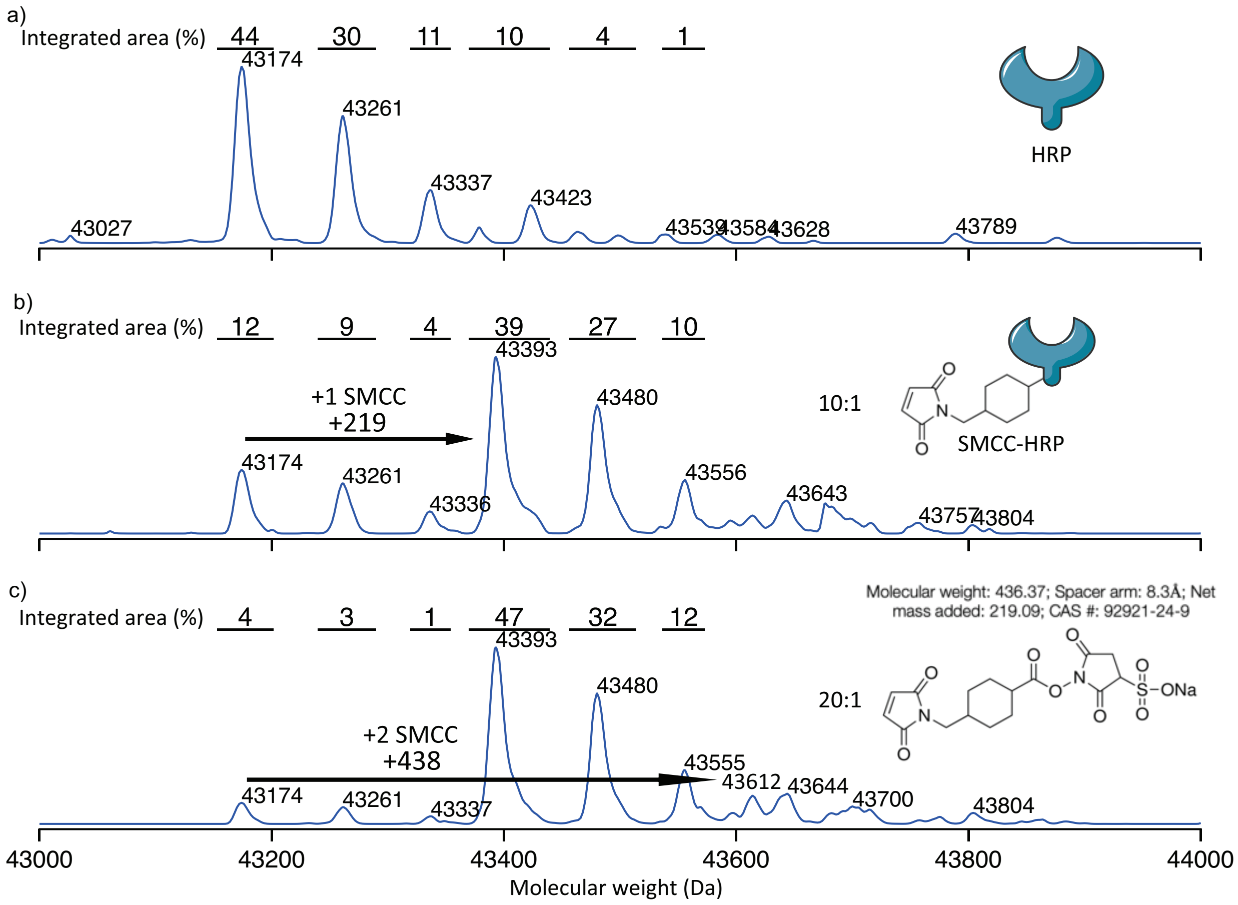


**Figure S1**. SMCC-HRP conjugation ratio optimization. Quantitative time-of-flight mass spectrometry (QTOF-MS) of HRP alone (a), SMCC-HRP reacted at a 10:1 ratio (b), and SMCC-HRP reacted at a 20:1 ratio (c) shows a ratio-dependent degree of maleimide functionalization. Though poly-dispersed, the primary mass of HRP alone by QTOF-MS reads as 43147 Da as seen in (a). According to the manufacturer, the mass added for one maleimide addition to HRP corresponds to about 219 Da, as seen by the appearance of the 44393 Da peak in (b). Similar additions of 219 Da to the other peaks present in (a) can be seen in (b) as well. Though (c) reveals 20:1 SMCC:HRP leads to a higher mono-functionalization than 10:1 as indicated by percent integrated area, it also exhibits higher dual-functionalization in the appearance of the 43612 Da peak. To minimize the potential of multiple maleimide additions reducing enzymatic activity, 10:1 SMCC:HRP was chosen for SWCNT conjugation.

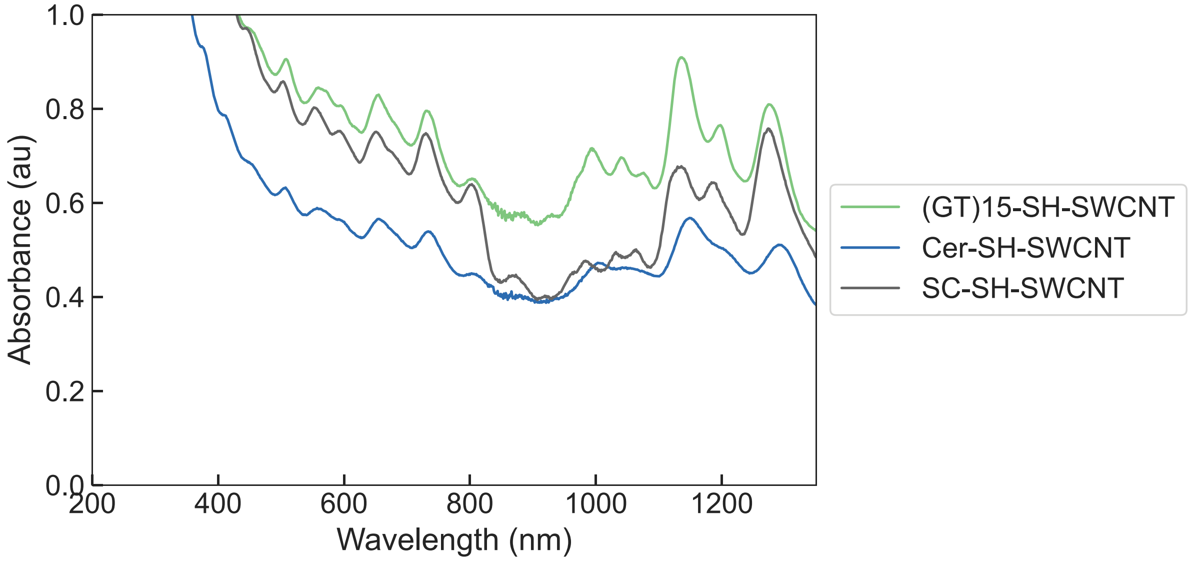


**Figure S2**. UV-Vis-IR spectra of SH-SWCNTs after probe tip sonication and centrifugation. After unsuspended SWCNT aggregates were removed by centrifugation, absorbance spectra were taken for each sample. All samples show expected SWCNT absorbance peaks in the nIR range (850 to 1350 nm), indicating probe tip sonication minimally interfered with the SWCNT $sp^2$ lattice. Each sample was diluted to normalize their absorbance to the linear range of Beer-Lambert's Law between 0 and 1 for accurate [SWCNT] calculation. The $(GT)_{15}$- and Cer-SH-SWCNTs were diluted 20X for absorbance measurements, while the SC-SH-SWCNTs were diluted 2X. The corresponding absorbances at 632 nm lead to measured [SWNCT] of 20.97, 14.65, and 19.44 mg $L^{-1}$, respectively and 419.49, 292.99, and 38.88 mg $L^{-1}$ when multiplied by their dilution factors.

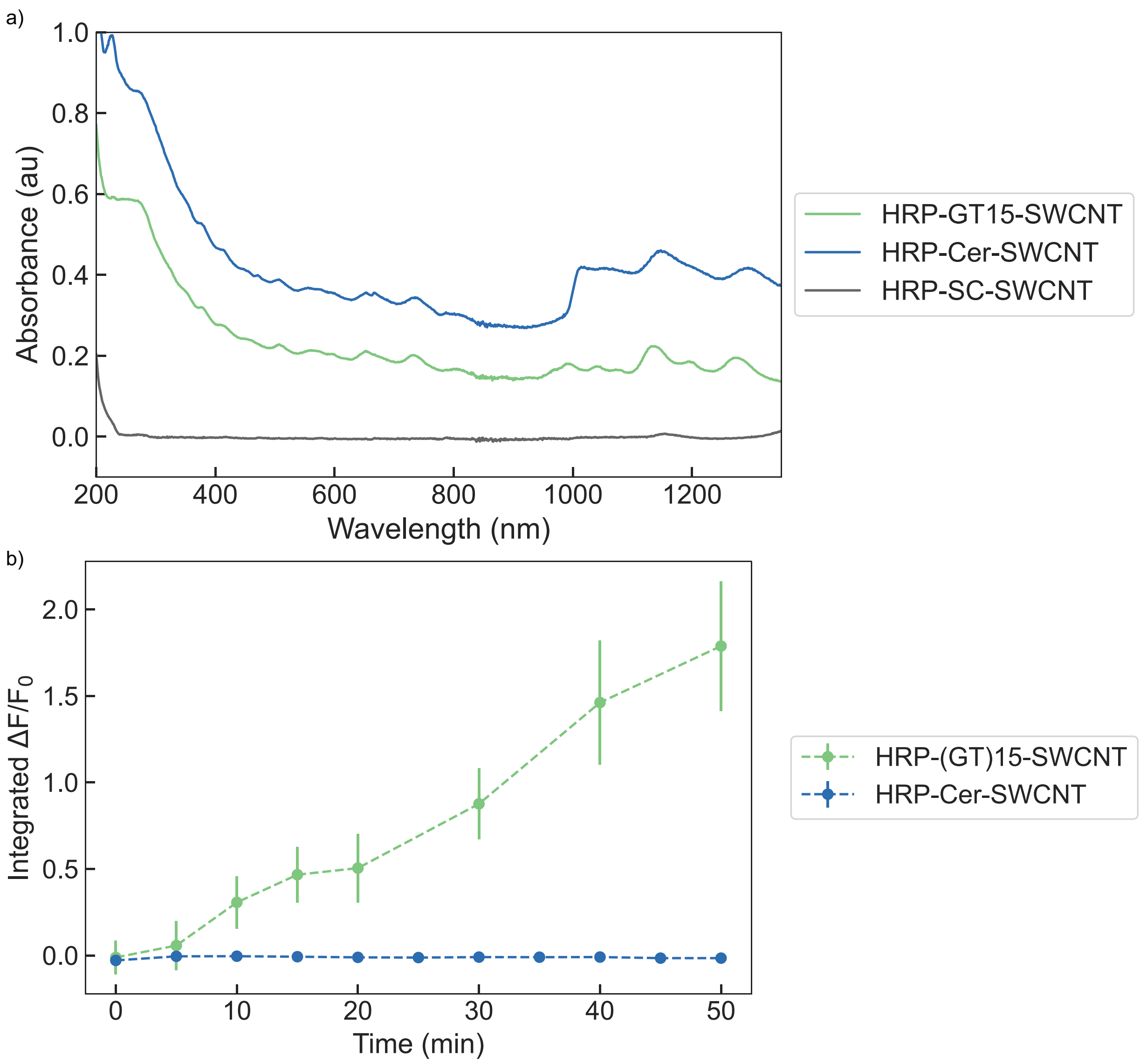


**Figure S3**. HRP-(GT)$_{15}$-SWCNT compared to other HRP-SWCNTs. (a) UV-Vis-IR spectra show loss of colloidal stability for HRP-SC-SWCNTs after purification as shown by the complete loss of characteristic SWCNT absorbance compared to (GT)$_{15}$- and Cer-coated HRP-SWCNTs. (b) The addition of 2941 µM of $H_2O_2$ to HRP-Cer-SWCNTs induces minimal fluorescence intensity change compared to the addition of the same concentration of $H_2O_2$ to HRP-(GT)$_{15}$-SWCNTs.

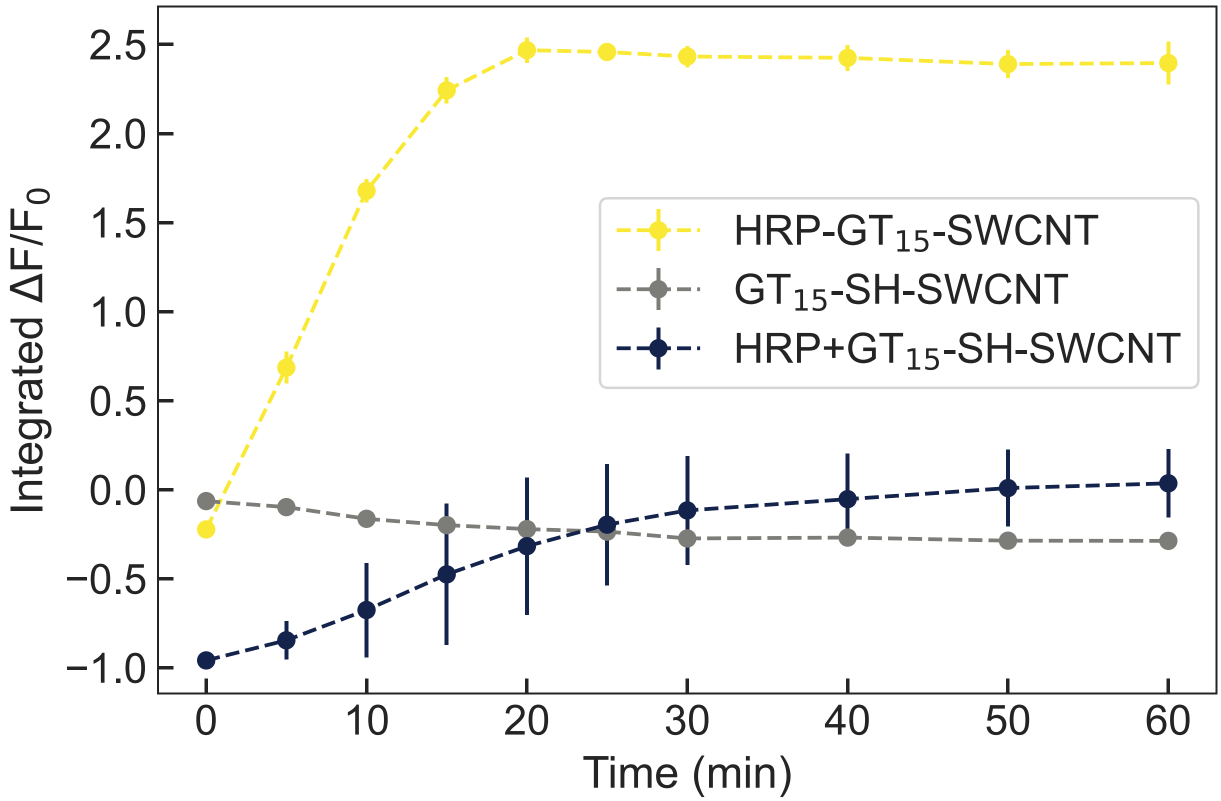


**Figure S4**. HRP-(GT)$_{15}$-SWCNT compared to starting material controls. Upon the addition of 29.4 mM of $H_2O_2$, (GT)$_{15}$-SH-SWCNT alone shows a slight turn-off response, replicating results from literature where $H_2O_2$ in proximity to the SWCNT surface quenches fluorescence.[31] For the addition of the same concentration of $H_2O_2$ to a solution of HRP mixed with (GT)$_{15}$-SH-SWCNT, a strong turn-off response is immediately observed followed by a gradual increase in fluorescence back to the original baseline intensity. This response could be attributed to the consumption of $H_2O_2$ by free HRP in the solution, reducing the quenching effect of $H_2O_2$ as the analyte is depleted. In contrast, HRP-(GT)$_{15}$-SWCNT exhibits a strong and stable turn-on fluorescence response to the same concentration of $H_2O_2$, highlighting the necessity of covalent HRP conjugation for this nanosensor's response.

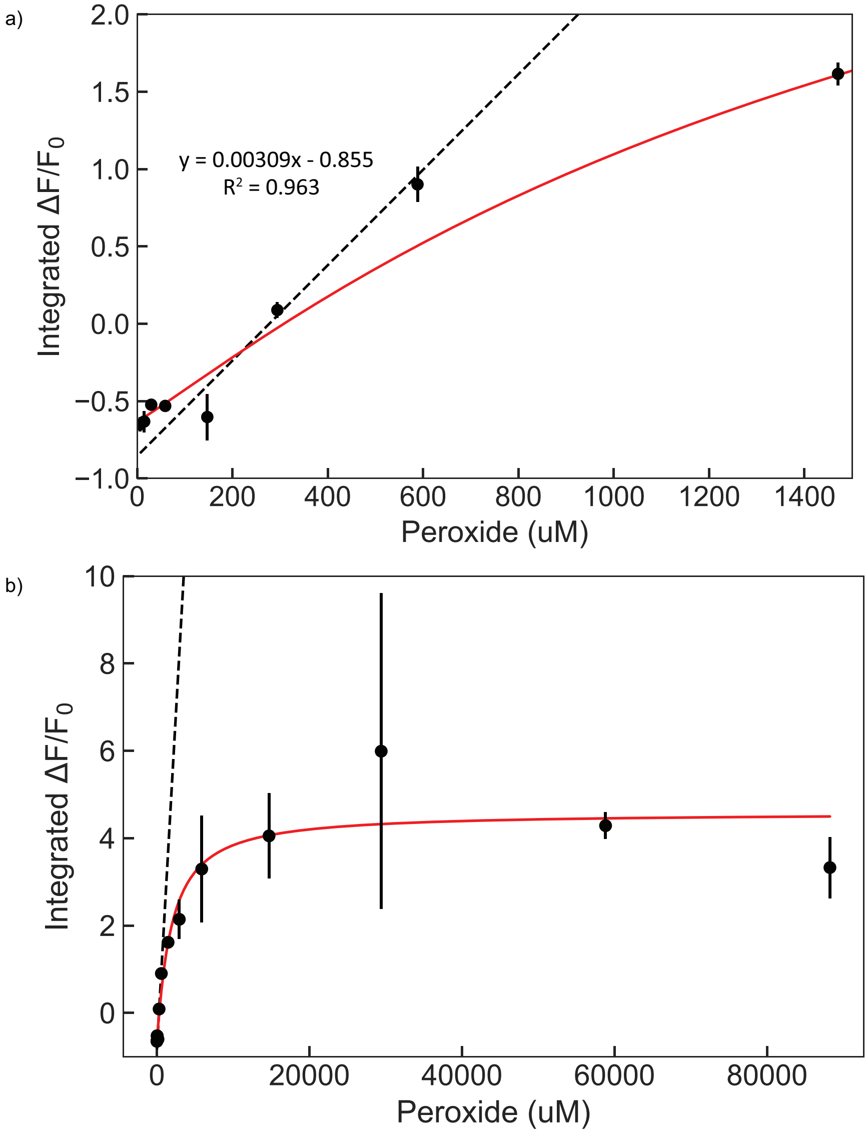


**Figure S5**. Linear range of HRP-(GT)$_{15}$-SWCNT nanosensor. Using weighted least squares regression, every possible set of 3 or more consecutive data points above the limit of detection was fit to a linear equation. (a) The region with the best linear fit as deemed by the highest $R^2$ value was between data points at 147.1 and 588.2 μM with $R^2 = 0.963$. All data points before the linear region and one data point after are included to demonstrate the deviation from linearity outside the identified range. (b) The plot of all data on a linear plot shows good agreement between the determined linear line and the visually observed region of linearity. Dotted black lines correspond to fitted linear equation and solid red lines correspond to fitted cooperative binding model.

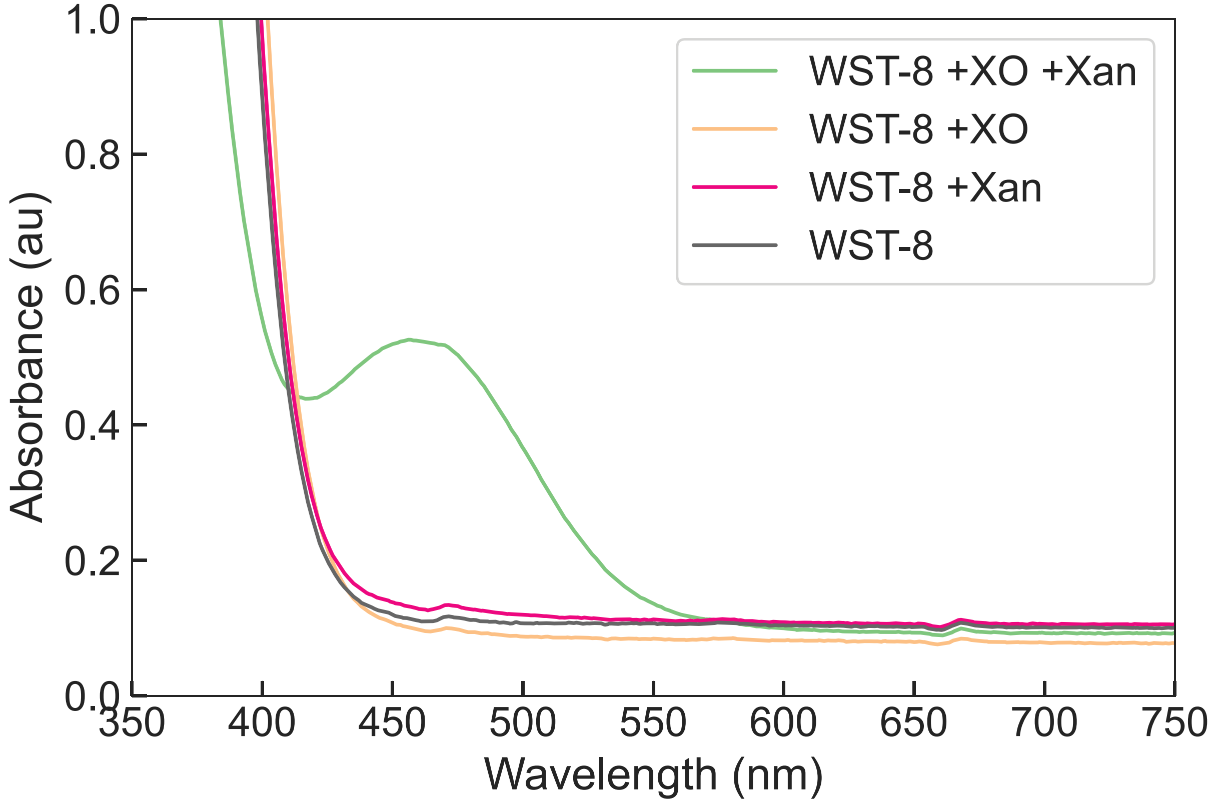


**Figure S6**. Validation of enzymatic superoxide generation with Xanthine/Xanthine Oxidase system and WST-8. Absorbance spectra for WST-8 is assessed alone, with Xanthine, with Xanthine Oxidase, and with both components. When superoxide is present, WST-8 is converted to WST-8 formazan, which exhibits an absorbance peak at 460 nm.[55] This peak at 460 nm is only evident when both Xanthine Oxidase and Xanthine are present, confirming the generation of superoxide under these conditions.
